# Biotechnology data analysis training with Jupyter Notebooks

**DOI:** 10.1101/2021.09.28.462133

**Authors:** Ulf W. Liebal, Rafael Schimassek, Iris Broderius, Nicole Maaßen, Alina Vogelgesang, Philipp Weyers, Lars M. Blank

**Affiliations:** iAMB - Institute of Applied Microbiology, ABBT, RWTH Aachen University, Worringerweg 1, 52074 Aachen, Germany; CLS – Center for Learning Services, RWTH Aachen University, Kackertstr. 15, 52072 Aachen, Germany

**Keywords:** Biotechnology, Data Analysis, Jupyter Notebooks, Python, Recombinant Expression, Systems Biology, Mathematical Model

## Abstract

Biotechnology has experienced innovations in analytics and data processing. As the volume of data and its complexity grows, new computational procedures for extracting information are developed. However, the rate of change outpaces the adaptation of biotechnology curricula, necessitating new teaching methodologies to equip biotechnologists with data analysis abilities. To simulate experimental data, we created a virtual organism simulator (*silvio*) by combining diverse cellular and sub-cellular microbial models. *silvio* was utilized to construct a computer-based instructional workflow with important steps during strain characterization and recombinant protein expression. The instructional workflow is provided as a Jupyter Notebook with comprehensive explanatory text of biotechnological facts and experiment simulations using *silvio* tools. The students conduct data analysis in Python or Excel. This instructional workflow was separately implemented in two distance courses for Master’s students in biology and biotechnology. The concept of using virtual organism simulations that generate coherent results across different experiments can be used to construct consistent and motivating case studies for biotechnological data literacy.

## Data Literacy in Biotechnology

Biotechnology is increasingly generating vast amounts of complex data that require advanced computational analysis (*1*). The experiments include, among others, multi-omics investigations (*2*), high-throughput techniques, or multiplexed experiments (*3*) and computational modeling approaches demand advanced data skills (*4*). In parallel, data science provides new tools to facilitate data analysis in the form of accessible machine learning tools, for example, via the Python SciKit learn library (*5*) and data analysis environments like Jupyter Notebooks (*6*), Galaxy (*7*), KBase (*8*), or KNIME (*9*). Particularly data analysis with Jupyter Notebooks are becoming popular (*10*) and have been used to guide metabolomic data analysis (*11*), metabolic engineering (*12, 13*), or gene expression (*14*). However, the developments of analysis have rarely managed to receive adequate attention in the curriculum of biotechnology.

Increasing the share of data analysis and bioinformatics in the biology and biotechnology curricula are excellent means for developing critical 21^st^-century skills by fostering inquiry-based interdisciplinary learning (*15*). Motivating and activating students is particularly important because the topic can deviate from many students’ prime interests and skills. Several reports discuss problem-based learning and flipped classrooms in computational biology (*15*–*18*). Also, Jupyter Notebooks are popular teaching resources in engineering (*19, 20*) and guidelines for their setup are available (*21, 22*). A particularly useful tool for motivation can be *gamification* and *serious game* elements (*22, 23*), which contribute more strongly to positive self-assessment than exam results (*24*).

Several virtual organism simulations allow precise phenotypic simulations of microorganisms. Models of organisms are increasingly extending from the fundamental mathematical representation into programmatic environments (*25*). Python is particularly popular because of the extensive community that supplies and maintains easily accessible packages to support general tasks from machine learning up to specific biological solutions (*25*). An outstanding recent development of virtual organism simulations is *Vivarium* as a modular environment that enables multi-scale models for realistic simulations (*26*).

Here, we report about a computational data analysis course for biotechnology, including growth curve analysis and recombinant gene expression. The data simulation is based on a collection of models that are combined into a unifying Python environment. The curse is designed as a tutorial in a Jupyter Notebook and although the courses were conducted distantly, small student groups jointly worked on simulations and data analysis. An evaluation of the course was performed by anonymous feedback about prior knowledge, competence inventory and experience, and usability of the Jupyter Notebook.

## Software and Methods

### The Simulator of Virtual Organisms

The microbial phenotypes are simulated in Python with an ensemble of models connected with the *silvio* (**si**mu**l**ator of **vi**rtual **o**rganisms) framework. *silvio* generates surrogate molecular and microbiological data without the attempt to reproduce real phenotypes. Instead, at each start of *silvio*, a new virtual organism is initiated with unique parameters to generate realistic data regarding type, structure, and complexity. The data is simulated by models of the associated experiments, including mechanistic models, *i*.*e*., growth laws, and statistical models, *i*.*e*., random forest for gene expression prediction. Figure 1 provides an overview of the architecture of *silvio*.

**Figure 1:**
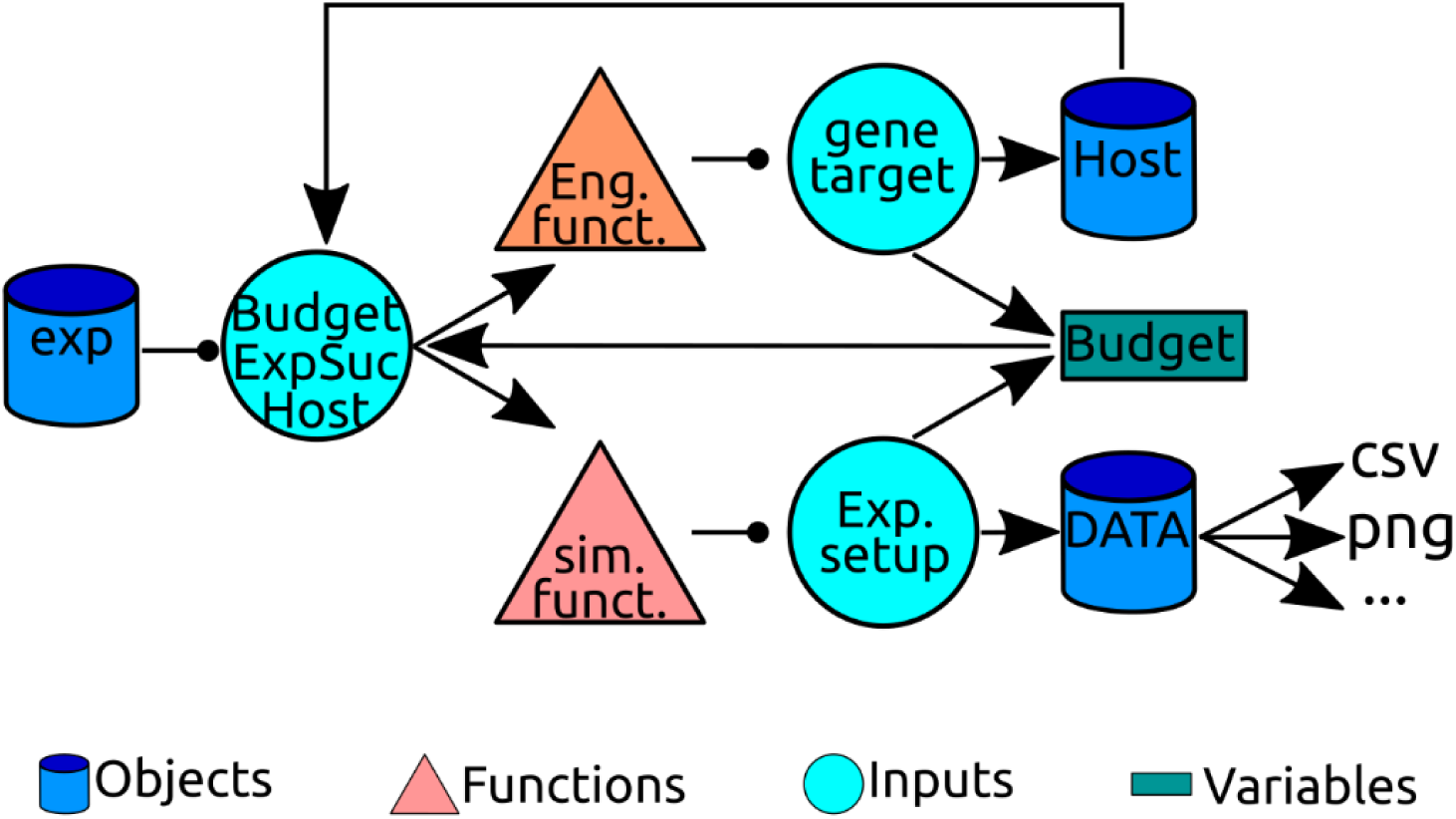
silvio code logic. A global experiment object (exp) is the starting point of functions for host engineering and genetic manipulation (Eng. funct.) and for analytical experiments (Exp. funct.). Both function require input about the host for the experiment (Host), available money (Budget) and experiment success probability (ExpSuc). The host organism is defined as an object with user-defined properties (strain type) and randomly initiated variables (optimal temperature, maximum biomass). Simulation data is stored in data objects with associated functions for post-processing.

*silvio* is organized in a parent object (*exp*) from which all experiments and genetic manipulation is performed. Two types of functions can be distinguished, one for changing host parameters and another for performing analytical experiments. The host object class contains all biological relevant information, for example, the host strain (*e*.*g*., *E. coli, P. putida*), the maximum biomass of cultivation (strain-specific, 30-145 g_CDW_/L), the optimal temperature (25-40° C), the optimal primer length for cloning (16-28 nucleotides), a correction factor for the expression strength (30-50), as well as budget information. The budget represents money which has to be invested in the initial lab equipment and in each experiment. The budget represents a gamification element and provides the students with estimates for the material requirements and the importance of an experiment. Importantly, students realize that a massive random search is impossible and they have to make well founded decisions. The amount of investment in the initial lab equipment determines the probability of experiment failures. In addition to the budget resource, the time dimension is considered such that experimental functions are designed to last a few seconds. Host-changing functions alter the values stored in the host object. Cloning, the only current host-changing function, duplicates the host properties with changes in the genetic information. The data is stored in separate objects to facilitate processing, visualization, and export.

*silvio* contains several models to simulate biological processes which are measured or optimized during biotechnological strain engineering and include growth, the growth constant, DNA melting temperature, and gene expression (Table 1). A logistic growth model based on the Verhulst equation simulates the growth experiments to determine optimal growth temperature. The maximum biomass (*K*) is randomly initiated between 30-100 and 45-145 g_CDW_/l for *E. coli* and *P. putida*, respectively. The temperature dependence of the growth constant (*r*) is calculated via a normal distribution function with a solution of 1 at the optimal temperature (corresponding to mean value μ) with variance σ=5. The optimal temperature (μ in normal distribution) is randomly initiated between 25-40° C.

**Table 1:** Models in silvio for biological processes.

| Process | Model | Details |
| --- | --- | --- |
| Growth | Verhulst logistic model<br>$P(t) = \frac{K}{1 + \left(\frac{K - P_0}{P_0}\right) e^{-rt}}$ | $P(t)$: biomass concentration<br>$K$: max. biomass, random<br>$P_0 = 0.1 \text{ g}_{\text{CDW}}/\text{L}$, initial biomass<br>$r$: growth constant, temperature dependent |
| Growth constant | Normal distribution | $\mu$: mean, optimal temperature, random<br>$\sigma = 5$: variance |
| DNA melting | $T_m = 2(A + T) + 4(C + G)$<br>$T_m = 64.9 + 0.41 \cdot GC(\%) - \frac{600}{N_{\text{NT}}}$ | $A, C, G, T$: number of bases<br>$GC(\%)$: GC content in %<br>$N_{\text{NT}}$: |
| Gene Expression | Random-Forest Regression | *E. coli*, *P. taiwanensis* promoter library, (Neves *et al.*, manuscript) |

Empirical formulas are used to calculate the DNA melting temperature and the gene expression strength. The optimal primer length is randomly initiated between 16-28 nucleotides (*27*). Two equations are used to calculate the optimal melting temperature, the first equation for primers <25 nucleotides, and the second equation for larger primers (Table 1). Gene expression is predicted based on the promoter sequence with 40 nucleotides (nt) upstream of the open reading frame. The regression is performed by a random forest machine learning module trained with measurements of a σ^70^ dependent synthetic promoter library expressed in *E. coli* and *P. taiwanensis* (*14*)(Neves *et al*., manuscript in preparation). The *silvio* simulation environment is available at https://git.rwth-aachen.de/ulf.liebal/silvio and the Jupyter Notebook is available at https://github.com/uliebal/RecExpSim/. A link to Binder is provided, which allows Cloud testing of the functions in a browser.

## General teaching setup

An instructional workflow was developed to convey biotechnological principles of recombinant expression within a modern data analysis environment. To increase motivation for the students, we developed a story about a fictional biotech company that aims to engineer a strain for vaccine production of a coronavirus protein subunit including a competition for the highest production rates. The steps included in the simulation are (1) choice of the host organism and its characterization, (2) design and cloning of a promoter sequence, and (3) measurement of the final expression rate. The final rates depend only on the promoter sequence and the precision by which the students measure host growth properties. The principle learning outcomes are described in Table 2 and include biotechnological knowledge as well as computer science procedures. During the simulation, different experiments are performed by using predefined functions appropriately. Each experiment requires resources: the money-budget is limited and some experiments are time-consuming to stimulate economic and effective use of experiments (here function calls from *silvio*).

**Table 2:** Principle learning outcomes of the Recombinant Expression Simulation Jupyter Notebook workflow.

|  |  |
| --- | --- |
| Informative | Growth and biomass are strain specific<br>Unknown factors impact cloning efficiencies<br>Promoter architecture of -35 and -10 boxes |
| Performative | Growth curve analysis with linear optimization<br>Calculation of DNA melting temperature<br>Variable assignment in Python |

The teaching unit was conducted as part of an otherwise lab-based biotech training in genetic manipulation and fermentation. The students were in their first year of Master curricula in biology and biotechnology at the RWTH Aachen University and the Westfälische University of Applied Science, respectively. The course was conducted each as Zoom-Meetings with an initial introduction and a walk-through of the simulation, including the solution to all steps (45 minutes). Following, participants were allotted to two-person groups in breakout rooms to work autonomously for ∼2 h on solving the simulation with regular (∼20 minutes) contact with a supervisor. The participants managed to test at least one and some even up to four promoter sequences for vaccine expression. In the last ∼20 minutes, all participants joined a conclusion round, during which the statistical relationship of promoter sequence and expression was examined, the production rates were compared and the group best production was honored.

### Data analysis in recombinant expression

The recombinant engineering workflow is simulated with four selected steps and data analysis tasks (see Table 3). The users start by choosing a host bacterium, predefined as either *E. coli* or *P. putida*. Both are popular bacterial hosts and related to important pathogens. The host choice affects the possible maximum biomass concentration and the predictor for the promoter activity but does not affect the final rate (see Methods for details). Along with the host, the user also decides how much money is invested in the laboratory equipment. An increasing investment correlates hyperbolically with decreasing experiment failures. The students started with 10,000 EUR starting capital, of which 10-20% are optimally used for the equipment.

**Table 3:** Computational steps of the Jupyter Notebook workflow. Each step involves a set of different activities and challenges.

| Step | Activity | Challenge |
| --- | --- | --- |
| 1. Host choice | • Read general introduction | • Python variable assignment |
| 2. Host characterization | • Analyze Excel data<br>• Estimate growth curve parameters | • Variance in biomass concentration<br>• Growth experiment fails |
| 3. Promoter design | • Learn importance of promoter boxes<br>• Calculate DNA melting temperature<br>• Derive primer complementary to promoter | • Optimal primer length unknown |
| 4. Evaluation | • Judge effectiveness of promoter<br>• Test impact of GC-content on expression | • Multiple rounds of promoter design |

### Host characterization

In the second step, the user identifies the optimal growth temperature and the associated growth rate and biomass. The temperature is randomly initiated between 25-40° C, the growth rate at the optimal temperature equals 1/h and biomass is randomly initiated (see Methods) in each simulation. The user inputs a vector with the test temperatures as an argument to the experiment function from *silvio* (*simulate_growth* in the experiment class) to identify the optimal temperature. The experiment for each temperature costs 100 EUR and takes ∼3 s simulation time. With a low probability of ∼10%, depending on investment in equipment, the growth fails and remains at the inoculum level. The outcome of this experiment is a CSV-file and related to the format of the *GrowthProfiler* (*Enzyscreen BV*). In our courses, the students could choose subsequent data analysis in *Excel* or *Python*. In *Excel*, the data has to be correctly imported, followed by logarithmic operation and visualization. The experimental column with the highest slope is then be subjected to linear regression within the linear regime of the logarithmic data to determine the growth rate and the average biomass during the stationary phase in the original data. To encourage Python-based analysis, we provided the necessary Python code for the data analysis but disordered the lines of code (Parsons puzzle). The students have to understand and rearrange the script. Two Python blocks exist: for visualization (Figure 2) and linear regression to extract growth rate and biomass. Analogously to the *Excel* procedure, the user identifies the optimal temperature along with growth rate and biomass.

**Figure 2:**
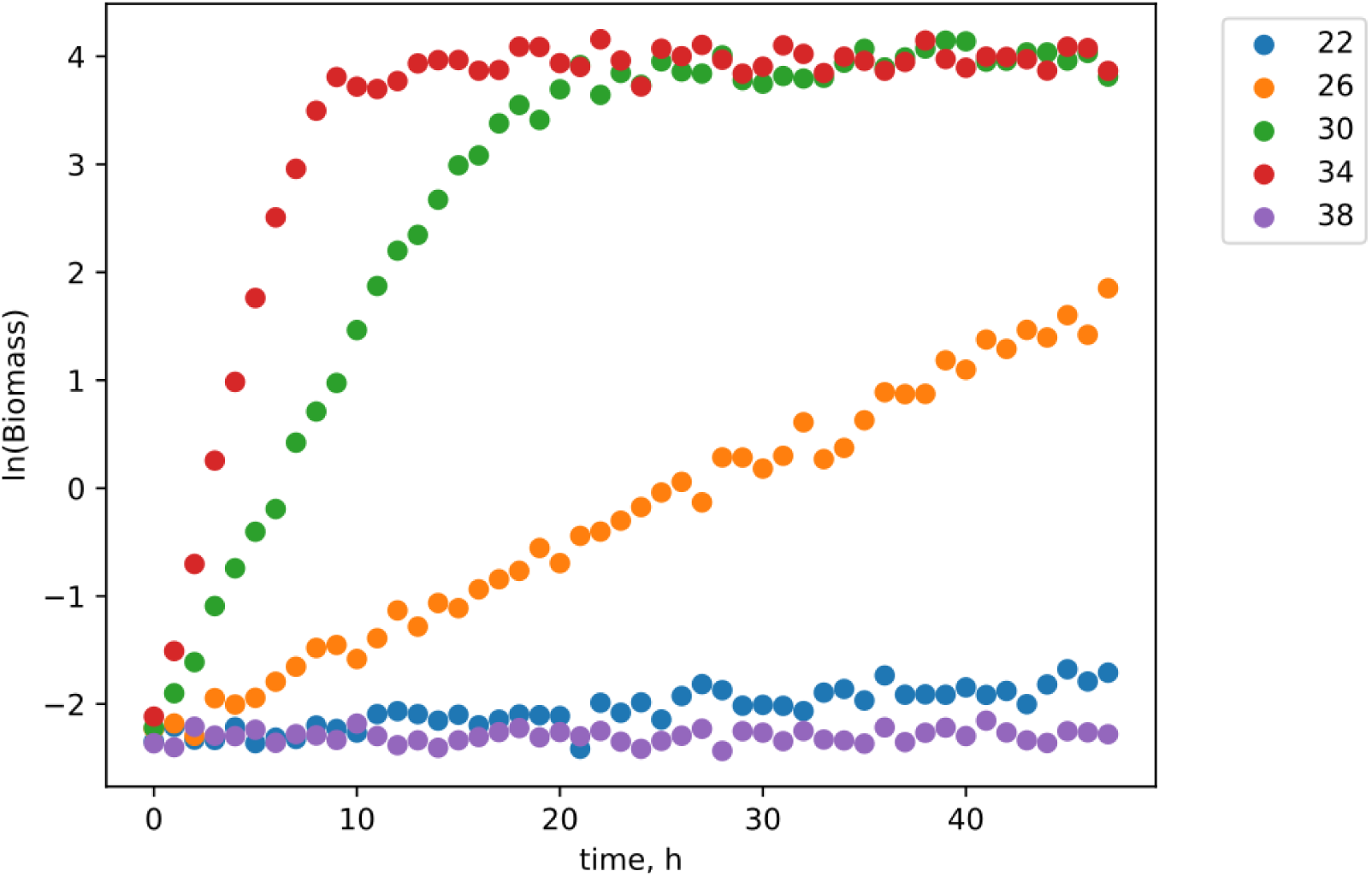
Example experiments (22-38° C) to determine optimal growth temperature. Five temperatures separated by 4° C were measured, which takes about 15 seconds of simulation time. The experiment at 34° C displayed the most robust linear growth increase until ∼7 h. The experiment at 38° C failed, with an expected growth rate comparable to the experiment at 30-34° C. Very slow growth is measured at the suboptimal temperature of 22° C.

### Promoter construction and cloning

After characterizing the host organism, the user is tasked to identify a suitable promoter sequence and cloning it into the host. The key learning objectives are knowledge about the structure of a standard sigma70 promoter, including boxes that control prokaryotic gene expression, computation of DNA melting temperature, and appreciation of the fickleness of cloning. The users are provided with a 40 nt promoter sequence reference:

GCCCA**XXXXXX**AXGCXXXCXCGTXXXGG**XXXXXX**TGCACG,

The *X* represents positions which the user has to replace with standard nucleotides and the bold printed X represent biologically important –35 and –10 boxes, respectively. The optimal composition of the box positions is explained in the accompanying text and linked to the literature (*28*). After the students replaced the *X* with nucleotides, the primer has to be developed. The primers start with the first nucleotide of the promoter but the optimal primer length is unknown and randomly initiated between 16-28 nt. The user then calculates the melting temperature according to the basic formula given by the equation in Table 1. The cloning experiment has a higher error rate and identical configurations will lead to different results with relatively frequent cloning failures. The cloning process is time-consuming and weakly predictive and thus mirrors somewhat the process of learning of a beginner. The students are motivated to try several versions of promoters to improve expression.

### Simulation results and cross-connection

Following the successful design of promoters, the users measure vaccine expression values, examine correlations of GC content and expression and identify the most effective host. The learning objectives are that multiple promoters need to be constructed to find a good one and that the GC content is not predictive of expression strength. The users first measure the promoter strength of each promoter construct. This evaluation is based on a Random Forest model trained on promoter libraries as described in the Materials and Methods section (*14*). The final experiment (*simulate_vaccine_production* in the experiment class) reports the vaccine production only if correct values derived from the host characterization experiments are provided (optimal temperature, biomass, and growth rate are within 10% of the correct values). As each group will only have tested 1-4 promoter variants, combining the results across the groups can lead to further insight. In our courses, we investigated correlations between GC content and vaccine expression strength for all tested promoters. An online audience response system (ARS NOVA) showed a blank plot with GC-content on the x-axis and expression strength on the y-axis with an underlying Hot-map feedback function. The students clicked on the corresponding value pairs of their tested promoters, which were stored and resulted in the plot in Figure 3. The students could appreciate that the combination of all data showed no direct correlation between GC content and vaccine expression. (ARS NOVA has meanwhile been converted into a commercial product and the Hot-map function was discontinued altogether.)

**Figure 3:**
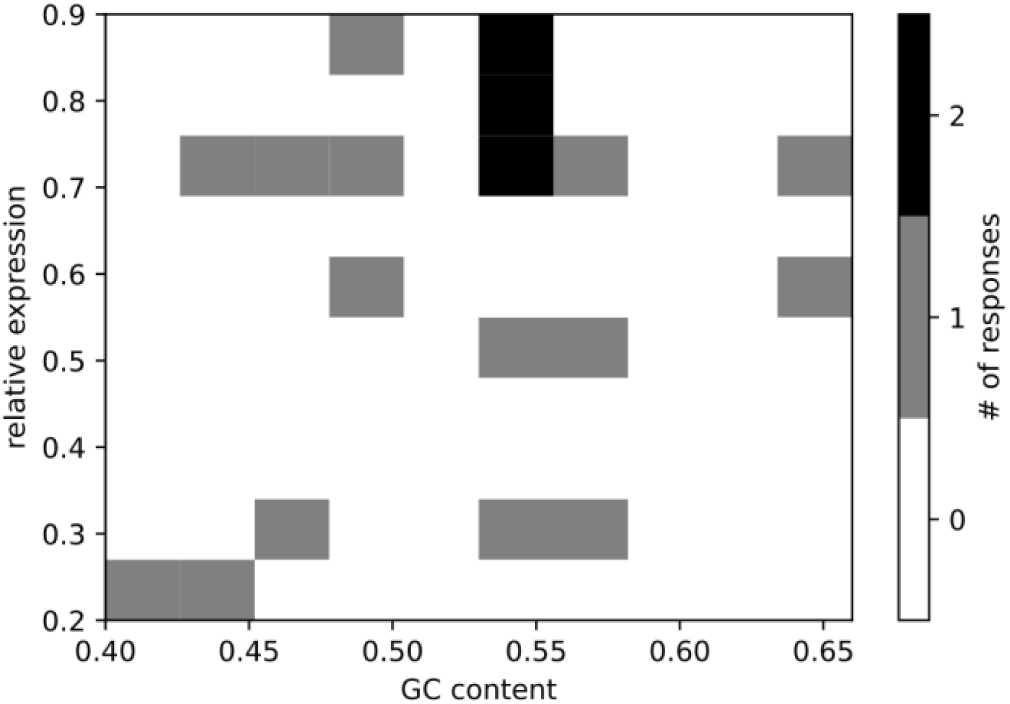
Joint simulation result of all groups with the relationship of GC-content to expression strength. It shows the potential to combine results from all groups to arrive at new knowledge when the effort of all groups is integrated.

## Course Feedback Survey

The course was supported with an evaluation to estimate the didactic benefit of the teaching concept. Evaluation is a critical component in optimizing and improving study programs and circumstances, as well as teaching in general. In all aspects of quality development processes, feedback is an important tool: whether as an inventory at the start of a project, selective feedback during the process, or as a final review of the effectiveness of measures - the achievement of objectives can be tracked at all stages. Established questionnaires in this study were used to obtain comparable, accurate and trustworthy data. The teaching evaluation took place at the end of the course, between the 21^th^ of April and the 25^th^ of May 2021, via an online questionnaire implemented on the platform *SoSciSurvey*. The results were evaluated with the statistics programme *R*. In total, 16 students took part in the survey. Due to the small sample size, a statistically reliable analysis of the results is not guaranteed.

### Survey instruments for teaching evaluation

To evaluate the educational benefit of the Jupyter Notebook based course, we assessed the prior knowledge, competence experience and competence gains as well as the increase in the topic related interest. We were also interested in how the course participants received the user-friendliness of the Jupyter Notebook and how they rated the course overall. Competence experience was assessed using a KIM questionnaire (Short scale of intrinsic motivation) (*29*). The questionnaire contained twelve perception-related statements (*e*.*g*., joyfulness, satisfaction, distress) and students rated the extent of agreement on a scale of 1 (full oposition) to 5 (full agreement). Competence gains were explored using the Graz Evaluation Model of Competence Acquisition (GEKO) for higher education courses (*30*). The GEKO contains a list of six different areas, for example “*the acquisition of theoretical, subject-related knowledge (e*.*g*., *theories and their contexts)*” or “*the ability to work in a team (e*.*g*., *coordination of cooperation with fellow students, division of tasks)*”, reported, again on the 1-5 scale. This scale of assessment was also used to evaluate increase in topic interest and the prior knowledge of students (novice to expert). A newly designed question section assessed the usability: experienced independence, the user-friendliness of the interface, the topic-related knowledge, and the complementarity to the parallel lecture (1-5 scale). The survey ended with the students’ overall evaluation of the course by seven self-developed questions (1-5 scale).

### Evaluation of the Survey

The evaluation revealed that, on average the students stated that they had rather little prior knowledge before attending the course (*M* = 1.75, *SD* = 1.13), and that the competence gain was rated relatively high (*M* = 3.66, *SD* = 0.56). The course also succeeded to raise interest in the topics and competencies covered (*M* = 3.88, *SD* = 1.02). In particular the acquisition of competences in teamwork received the highest rating and testifies the effectiveness of digital collaborations (*M* = 3.94, *SD* = 1.12). The evaluation of the KIM questionnaire showed that the course increased intrinsic motivation and the experience of competence on average (*M* = 3.37, *SD* = 0.49). Regarding the usability, the evaluation revealed that students were initially concerned about struggling with the computational tasks, but in the end, they found it enjoyable and the simulations were rated overall as user-friendly (*M* = 3.67, *SD* = 0.55). The total score rating of the course also indicates that the students enjoyed the course (*M*= 3.58, *SD* = 0.43). (Figure 4). Overall, the feedback demonstrated that the Jupyter Notebook based course led to a generally positive learning experience.

**Figure 4:**
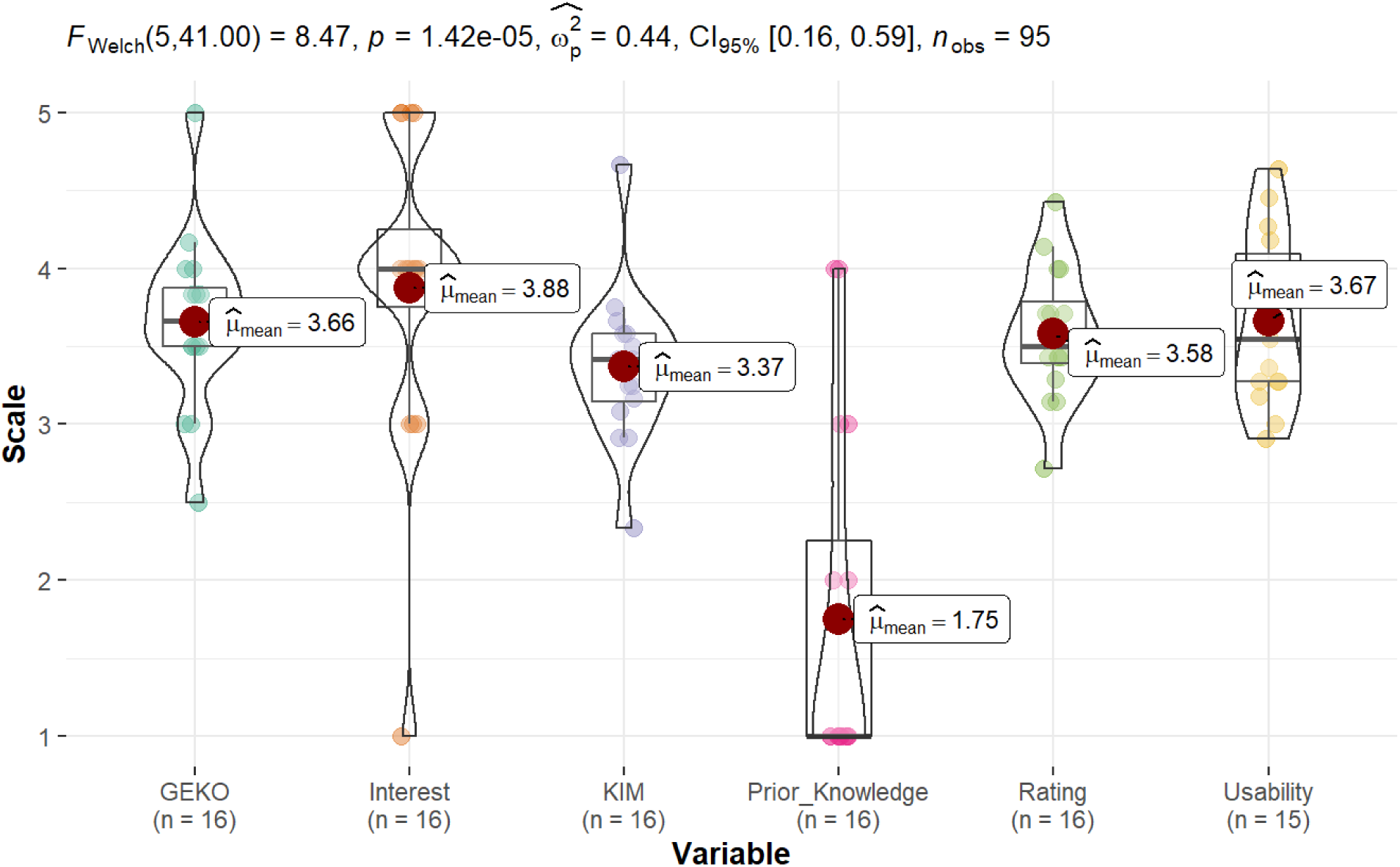
Distributions of the total scores of the scales regarding competence acquisition (GEKO), the students’ assessments of the extent to which their interest in the topics was increased by the course (Interest), the perceived competence experience (KIM), how the students assess their prior knowledge before attending the event (Prior_Knowledge), the Overall Evaluation of the Event (Rating) and the Usability of the Jupyter Notebook.

## Conclusions

We have developed a model ensemble to simulate biotechnological relevant experiments for microorganisms (*silvio*). These experiments were combined in an instructional Jupyter Notebook to teach selected data analysis steps in a recombinant gene expression project. By adding new models to *silvio*, new biotechnological experiments can be simulated and converted into instructional, motivating data analyses projects in the form of Jupyter Notebooks. Our approach decouples the computational data analysis from often lengthy, expensive, and sometimes unavailable experiments. We stress that our simulations are not meant to replace the physical experience of conducting the experiments in the laboratory but rather to support analytical lectures by providing practice in data handling and evaluation. Finally, our survey documented the positive learning experience by the students.

